# Intrinsic Activity Develops Along a Sensorimotor-Association Cortical Axis in Youth

**DOI:** 10.1101/2022.08.15.503994

**Authors:** Valerie J. Sydnor, Bart Larsen, Jakob Seidlitz, Azeez Adebimpe, Aaron Alexander-Bloch, Dani S. Bassett, Maxwell A. Bertolero, Matthew Cieslak, Sydney Covitz, Yong Fan, Raquel E. Gur, Ruben C. Gur, Allyson P. Mackey, Tyler M. Moore, David R. Roalf, Russell T. Shinohara, Theodore D. Satterthwaite

**Author notes:** Correspondence: Theodore D. Satterthwaite.

## Abstract

Animal studies of neurodevelopmental plasticity have shown that intrinsic brain activity evolves from high amplitude and globally synchronized to suppressed and sparse as plasticity declines and the cortex matures. Leveraging resting-state functional MRI data from 1033 individuals (8-23 years), we reveal that this stereotyped refinement of intrinsic activity occurs during human development and provides evidence for a cortical gradient of neurodevelopmental plasticity during childhood and adolescence. Specifically, we demonstrate that declines in the amplitude of intrinsic activity are initiated heterochronously across regions, coupled to the maturation of a plasticity-restricting structural feature, and temporally staggered along a hierarchical sensorimotor-association axis from ages 8 to 18. Youth from disadvantaged environments exhibit reduced intrinsic activity in regions further up the sensorimotor-association axis, suggestive of a reduced level of plasticity in late-maturing cortices. Our results uncover a hierarchical axis of neurodevelopment and offer insight into the temporal sequence of protracted neurodevelopmental plasticity in humans.

## INTRODUCTION

Elucidating the spatiotemporal progression of developmental plasticity across the human cortex has implications for understanding healthy brain development as well as windows of developmental vulnerability and opportunity. Specifically, characterizing the temporal sequence of cortical plasticity is requisite for identifying biological mechanisms underlying normative developmental plasticity and the potential role of disrupted plasticity in youth-onset psychopathology^1^. Moreover, demarcating regionally-specific periods of enhanced and diminished malleability can reveal windows where both developmental insults and interventions may have a maximal impact on the brain^2^. Prior studies have therefore aimed to uncover the temporal sequence of neurodevelopmental change across the cortical mantle.

These studies have consistently shown that postnatal neurodevelopment is heterochronous, with sensory and motor cortices maturating earlier than association cortices; this temporal trend has been shown for cortical volume, thickness, connectivity, myelination, and cellular properties^3–12^. Critically, beyond this coarse division there exists marked temporal developmental variability that remains under-characterized. We recently proposed a unifying framework on the chronology of cortical development that contextualizes past reports of asynchronous maturation between sensorimotor and association cortices^2,13–16^ as two ends of a continuous axis of neurodevelopmental plasticity^1^. This framework posits that during childhood and adolescence, cortical plasticity progresses along the sensorimotor-association (S-A) axis: a dominant, hierarchical axis of human brain organization along which diverse neurobiological properties are patterned^1,17–20^.

In the present study, we aimed to empirically evaluate our hypothesis that plasticity unfolds along the S-A axis by studying the developmental refinement of intrinsically-generated (i.e., spontaneous or non-evoked) activity, a putative functional marker of local plasticity described in animal models. Studies of the developing murine sensory cortex have provided evidence that a potentiation of high amplitude, synchronized intrinsic activity characterizes early periods of heightened plasticity^15,21,22^. As plasticity is reduced and the cortex matures, intrinsic activity then evolves from strong and globally synchronized to suppressed and sparse, becoming more heterogeneously distributed in space and time in adult cortex^22–26^. This stereotyped refinement of intrinsic activity has been linked to maturational increases in inhibitory neurotransmission and cortical myelination—two plasticity-regulating processes that refine cortical circuit dynamics^21,26–30^. Accordingly, this stereotyped refinement of intrinsic activity provides an ongoing readout of local circuit plasticity, with more widespread and correlated high amplitude activity serving as a functional hallmark of immature, plastic cortices^15,21,31,32^. Importantly, intrinsic cortical activity can be studied non-invasively with resting-state functional MRI (fMRI), which provides an opportunity to characterize the temporal maturation of a plasticity signature in the human brain.

Simultaneous fMRI and electrophysiology or calcium recordings have demonstrated how low frequency fluctuations in the resting fMRI blood oxygen level dependent (BOLD) signal are coupled with changes in intrinsic neural activity patterns^33–38^. A greater level of intrinsic activity and more synchronized activity—activity characteristic of immature, plastic cortices—increases the amplitude of low frequency BOLD fluctuations. It has therefore been hypothesized that the amplitude of low frequency fluctuations^39^, or BOLD “fluctuation amplitude”, will be higher when cortical plasticity is enhanced^31,40^. Indeed, a recent landmark study found that casting of the upper extremity in non-injured humans—a deprivation intervention designed to induce somatosensory and motor cortex plasticity—increased BOLD fluctuation amplitude selectively within the corresponding topographic region of primary somatomotor cortex^41^.

Here we harness BOLD fluctuation amplitude to index spatially-localized, age-dependent changes in intrinsic cortical activity and test the overarching framework that developmental plasticity cascades hierarchically along the S-A axis in youth. Given evidence that intrinsic activity declines and desynchronizes as developmental plasticity is reduced, we predicted that the development of fluctuation amplitude would be characterized by heterochronous declines along the cortex’s S-A axis and would be linked to the maturation of plasticity-regulating biological features. Furthermore, in light of recent theories that cortical maturation is accelerated by environmental adversity^42–44^, we predicted that youth from disadvantaged neighborhoods would exhibit functional markers suggestive of lower cortical plasticity. As described below, our in vivo analysis of a signature of neurodevelopment plasticity illuminated by animal models reveals that the S-A axis captures not only the hierarchical layout of diverse cortical properties, but also the temporal patterning of human developmental programs.

## RESULTS

We studied how intrinsic activity is refined across the developing cortex in a sample of 1033 youth aged 8-23 years. Fluctuation amplitude, computed as the average power of low frequency (0.01-0.08 Hz) fluctuations in the time-varying fMRI signal, was used to index the overall level and coherence of intrinsic cortical activity. Greater and more synchronized neural activity increases the power of low frequency neural recordings^34,35,37,38^ and is understood to increase BOLD fluctuation amplitude^34,35^. To delineate maturational changes in BOLD fluctuation amplitude in individual cortical regions, we fit region-specific generalized additive models (GAMs) in which age was treated as a smooth term and modeled with thin plate regression splines; sex and in-scanner head motion were included as covariates. Each regional GAM estimates a smooth function (the model age fit) that describes the relationship between fluctuation amplitude and age; the first derivative of this smooth function, estimated with finite differences, represents the rate of change in fluctuation amplitude at a given developmental timepoint. We tested whether GAM-derived developmental effects provide support for our hierarchical neurodevelopmental plasticity framework; we also validated that effects were robust to controls for in-scanner motion, medication use, vascular effects, T2* signal strength, and cortical atlas. All code used to calculate fluctuation amplitude, fit regional GAMs, contextualize developmental effects, and perform sensitivity analyses is available along with detailed documentation for code implementation at https://pennlinc.github.io/spatiotemp_dev_plasticity.

### Age-dependent changes in intrinsic fMRI fluctuations vary across the cortex

Fluctuation amplitude significantly changed with age in the developmental window studied in nearly all cortical regions (*p*_FDR_ < 0.05 in 95% of regions), providing evidence that intrinsic cortical activity is refined from childhood to early adulthood. To provide insight into the overall magnitude and direction of regional age effects, we calculated the variance explained by age (partial *R*^*2*^; i.e., the effect magnitude) and the sign of the average derivative of the age fit (i.e., the effect direction). The magnitude and direction of age effects differed across the cortex (**Fig. 1A**), signifying there is variability in the maturation of fluctuation amplitude across the developing brain. Indeed, by visualizing age fits across regions we observed a cortical continuum of developmental trajectories ranging from large and prolonged decreases (light yellow fits in **Fig. 1B**) to inverted U-shaped curves (dark purple fits). Nearly all sensory regions showed continuous declines in fluctuation amplitude from early childhood through adolescence, as illustrated by the model fit for area V1 (**Fig. 1C**, top panel) which significantly decreased until age 18 years. This age fit is potentially indicative of a progressive reduction, sparsification, or decorrelation of non-evoked cortical activity with age. In contrast, in select cortical regions such as the midcingulate gyrus (**Fig. 1C**, middle panel), fluctuation amplitude only began to decline in later childhood or early adolescence. Finally, many regions in transmodal association cortex (e.g., the dorsolateral prefrontal cortex; **Fig. 1C**, bottom panel) displayed significant increases in fluctuation amplitude until early to mid-adolescence, followed by amplitude decreases. This inverted U-shaped trajectory suggests there is heightened, synchronized activity in transmodal cortices at the start of adolescence. The trajectories of regional age fits did not significantly differ between males and females, indicating that the timing of developmental change did not vary by sex in this age range (*p*_FDR_ > 0.05 for all age-by-sex interactions). These results establish that there is heterogeneity in the maturation of intrinsic fMRI fluctuations, with maturational trends broadly diverging between sensory and association cortices.

**Fig. 1.**
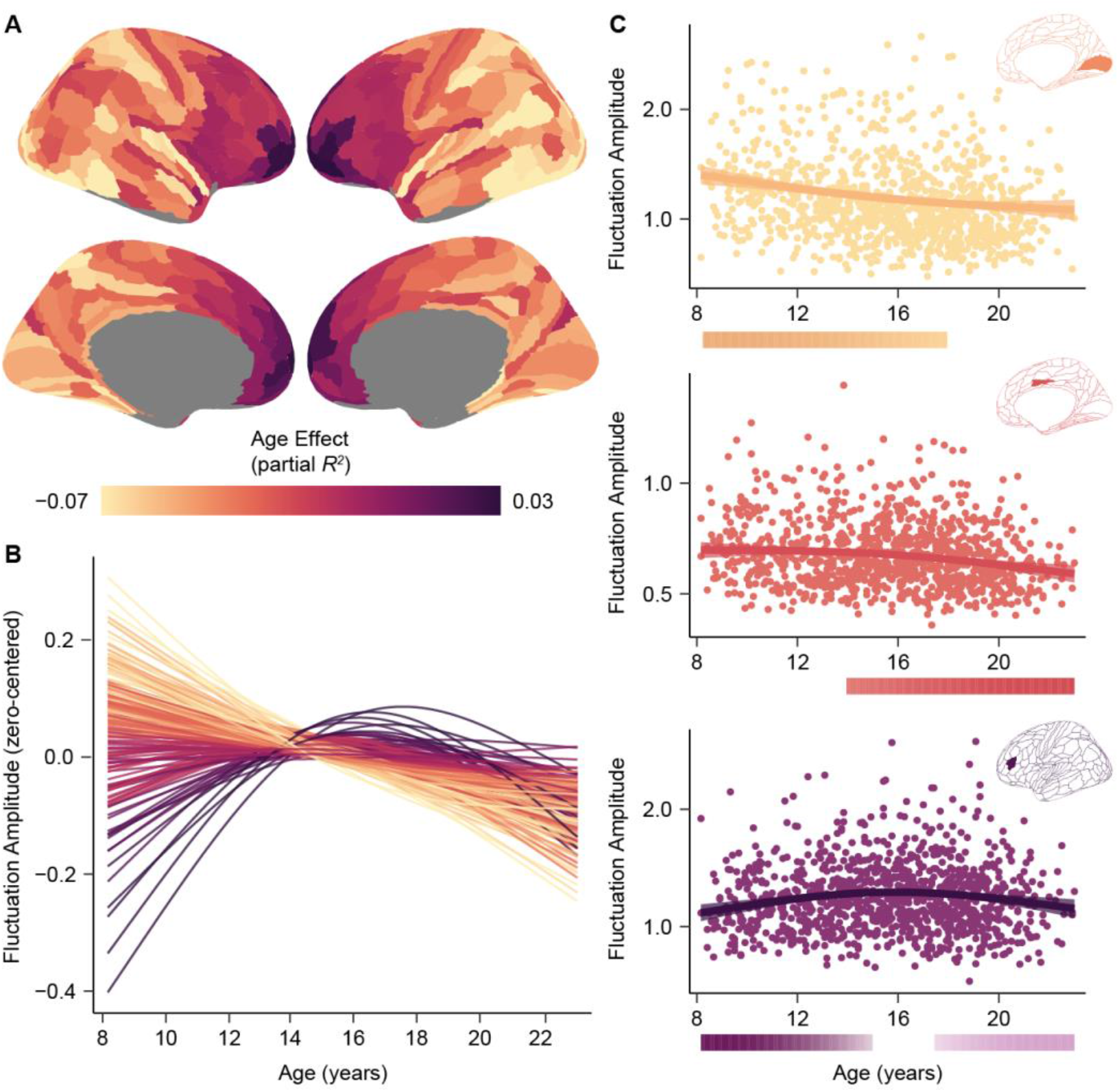
Age-related refinement of fluctuation amplitude varies across the cortex. **A)** The patterning of fluctuation amplitude age effects is displayed across the cortical surface. **B)** Fluctuation amplitude developmental trajectories (zero-centered) are shown for all left hemisphere cortical regions. Trajectories are colored by each region’s age effect using the color bar in panel A. **C**) Fluctuation amplitude developmental trajectories are shown overlaid on data from all participants for the primary visual cortex (area V1, yellow), the midcingulate gyrus (area p24pr, pink), and the dorsolateral prefrontal cortex (area IFSa, purple). The color bars below each regional plot depict the age window(s) wherein fluctuation amplitude significantly changed in that region, shaded by the rate of change.

### The development of intrinsic fMRI fluctuations is linked to the maturation of a key regulator of plasticity

Prior work in animal models has shown that intrinsic cortical activity develops from widespread, high amplitude, and synchronized to sparse and decorrelated as the cortex progresses from plastic to mature, signifying that age-dependent changes in the amplitude of BOLD fluctuations could, in part, reflect changes in cortical plasticity^31,41^. We therefore endeavored to understand whether the development of fluctuation amplitude is related to the maturation of intracortical myelination–a key regulator of cortical plasticity. In developing neural circuits, myelination constrains further axonal and dendritic plasticity and thus serves as a circuit stabilizer and plasticity-limiting factor^30,45^. Here, we tested whether fluctuation amplitude maturation was related to maturation of the T1w/T2w ratio, a structural MRI measure sensitive to cortical myelin content^46^.

To do so, we leveraged the recent work of Baum et al. (2022), who studied the development of T1w/T2w-indexed myelination in a large, independent sample of youth ages 8-21 years. These authors quantified myelination age effects (as partial *R*^*2*^) and demarcated the age of peak myelin growth within individual cortical regions. In comparing T1w/T2w ratio and fluctuation amplitude neurodevelopmental features, we unveiled substantial spatial and temporal correspondence between the refinement of these measures with age. Age-related changes (indexed by the signed partial *R*^*2*^) in these two putative *in vivo* indicators of plasticity were strongly inversely correlated across cortical regions (*r* = -0.67, *p*_spin_ < 0.001), with regions showing larger increases in myelin content from childhood to early adulthood also showing larger decreases in BOLD signal amplitude (**Fig. 2A-B)**. This finding accords with ample evidence of causal, bidirectional relationships between changes in neural activity patterns and changes in myelination^47–49^ and suggests a possible mechanistic link between microstructural refinement and reductions in intrinsic activity during brain development.

**Fig. 2.**
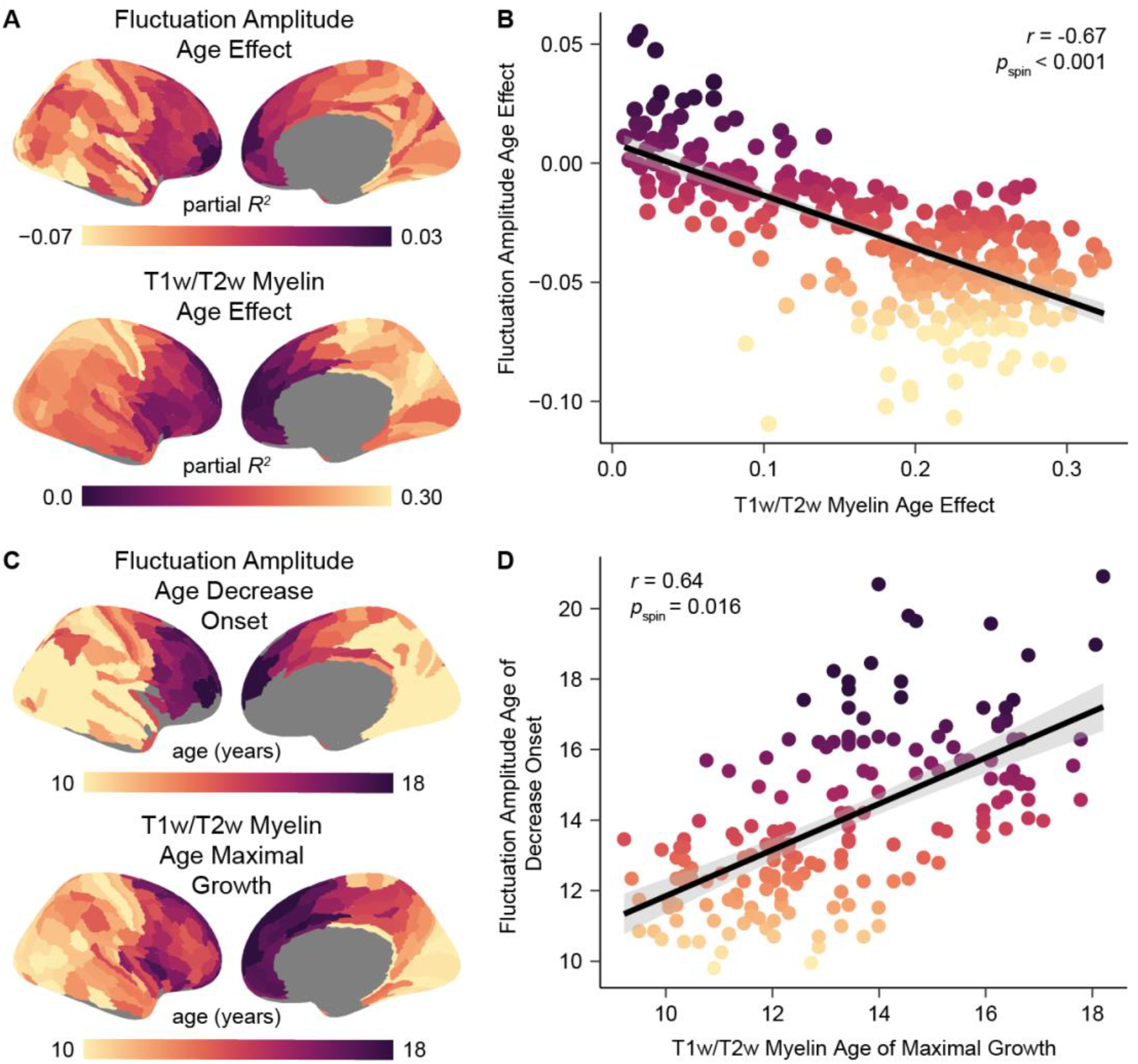
Development of fluctuation amplitude spatially and temporally parallels cortical myelin development. **A**) The cortical distribution of fluctuation amplitude age effects closely resembles the distribution of T1w/T2w ratio age effects, suggesting interdependent refinement of cortical function and microstructure in youth. Age effects are signed by the sign of the average derivative of the age smooth function. **B**) Regions that experienced larger declines in fluctuation amplitude during childhood and adolescence additionally underwent greater increases in intracortical T1w/T2w ratio in this developmental period. **C**) Maps depicting the age at which fluctuation amplitude began to decrease (earliest significant negative derivative of the age function) and the age of maximal T1w/T2w-indexed myelin growth (largest significant derivative of the age function) reveal temporal similarity in the development of these two measures in youth. **D**) Across regions, the age at which fluctuation amplitude began to significantly decrease is closely coupled to the age at which the T1w/T2w ratio showed a maximal rate of increase, providing evidence for temporal coordination between reductions in intrinsic activity and the development of cortical myeloarchitecture.

To further explore this link, we investigated whether there was a temporal relationship between increases in the T1w/T2w ratio and decreases in fluctuation amplitude during youth. We first quantified the age at which fluctuation amplitude began to significantly decrease in each region and found that initial decreases in fluctuation amplitude were staggered heterochronously across the cortex. Nearly half of regions (46%) showed a significant decrease in fluctuation amplitude at age 8, implying that BOLD amplitude within these regions likely begins to decline prior to the youngest age studied in our dataset. Across the rest of the cortex, however, fluctuation amplitude began to decline later in youth; in these cortices, a greater delay in the onset of fluctuation amplitude decline was associated with a later peak in the rate of T1w/T2w-indexed cortical myelination (*r* = 0.64, *p*_spin_ = 0.016; **Fig. 2C-D**). This association suggests that cortices that undergo maximal myelin growth at a later developmental stage also experience a temporally delayed decline in the amplitude of spontaneous fMRI activity. Notably, ages of fluctuation amplitude decrease onset and maximal T1w/T2w increase were not simply correlated but also showed a minimal temporal offset in years, indicating that they were closely coupled in time (average offset = 0.7 years; see also the best fit line for **Fig. 2D)**. Across regions, it was more common for the age of maximal T1w/T2w ratio increase to precede the age of fluctuation amplitude decline. Taken together, these results link age-related refinement of fluctuation amplitude to maturation of a main regulator of developmental plasticity.

### Spatiotemporal variability in developmental trajectories is captured by the sensorimotor-association axis

A primary goal of this work was to systematically assess whether the sequence of neurodevelopmental plasticity progresses hierarchically across the cortical mantle. Having observed that fluctuation amplitude development tightly paralleled development of a plasticity regulator and broadly diverged between sensorimotor and association cortices, we next sought to determine whether developmental patterns spatially conformed to the S-A axis^1^. The S-A axis is a prominent axis of cortical variation that is ordered from primary sensory and motor cortices, to modality-selective and multimodal cortices, then progressing to transmodal heteromodal and paralimbic cortices. This axis captures the concerted patterning of heterogeneous macrostructural, metabolic, cellular, molecular, transcriptomic, and electrophysiological properties across the cortical mantle^1,17–20,50^. Moreover, the S-A axis spatially coheres with the brain’s anatomical^50^, functional^20^, and evolutionary^51^ hierarchies, thus each cortical region’s rank in the axis reflects its relative position in a global cortical hierarchy.

We first examined whether inter-regional differences in the development of fluctuation amplitude reflected inter-regional differences in S-A axis rank. Region-wise age effects and S-A axis ranks were highly correlated (*r* = 0.54, *p*_spin_ = 0.002), with large negative age effects characterizing the S-A axis’s sensorimotor pole and smaller positive age effects distinguishing its association pole. We additionally observed continuous variation in the age at which fluctuation amplitude began to significantly decrease along this dominant organizational axis. When considering regions that showed an initial onset of decline within the age range studied as above, we found that fluctuation amplitude began to decline at a progressively later age in regions ranked higher in the S-A axis (*r* = 0.68, *p*_spin_ = 0.001). Hence, cortices at the top of the cortical hierarchy exhibit the smallest and latest-onset declines in the amplitude of intrinsic fMRI fluctuations during childhood and adolescence.

Following this initial analysis, we further probed the extent to which maturational trajectories differed as a function of S-A axis rank by mapping the principal spatial axis of fluctuation amplitude development. To accomplish this mapping, we performed a principal component analysis (PCA) on the age fits estimated by regional GAMs (**Fig. 1B**); this approach considers the entire fluctuation amplitude developmental trajectory rather than only one property of the age fit (e.g., the age at which it starts to decline). The first principal component from this PCA explained 87% of the variance in developmental profiles and can therefore be conceptualized as the principal axis of intrinsic activity development. This principal developmental axis closely resembled the S-A axis (**Fig. 3A**). Accordingly, regional loadings onto the principal developmental axis were very highly correlated with regional S-A axis ranks (*r* = 0.70, *p*_spin_ < 0.001; **Fig. 3B**), demonstrating that the vast majority of spatiotemporal variance in developmental profiles was explained by the S-A axis.

**Fig. 3.**
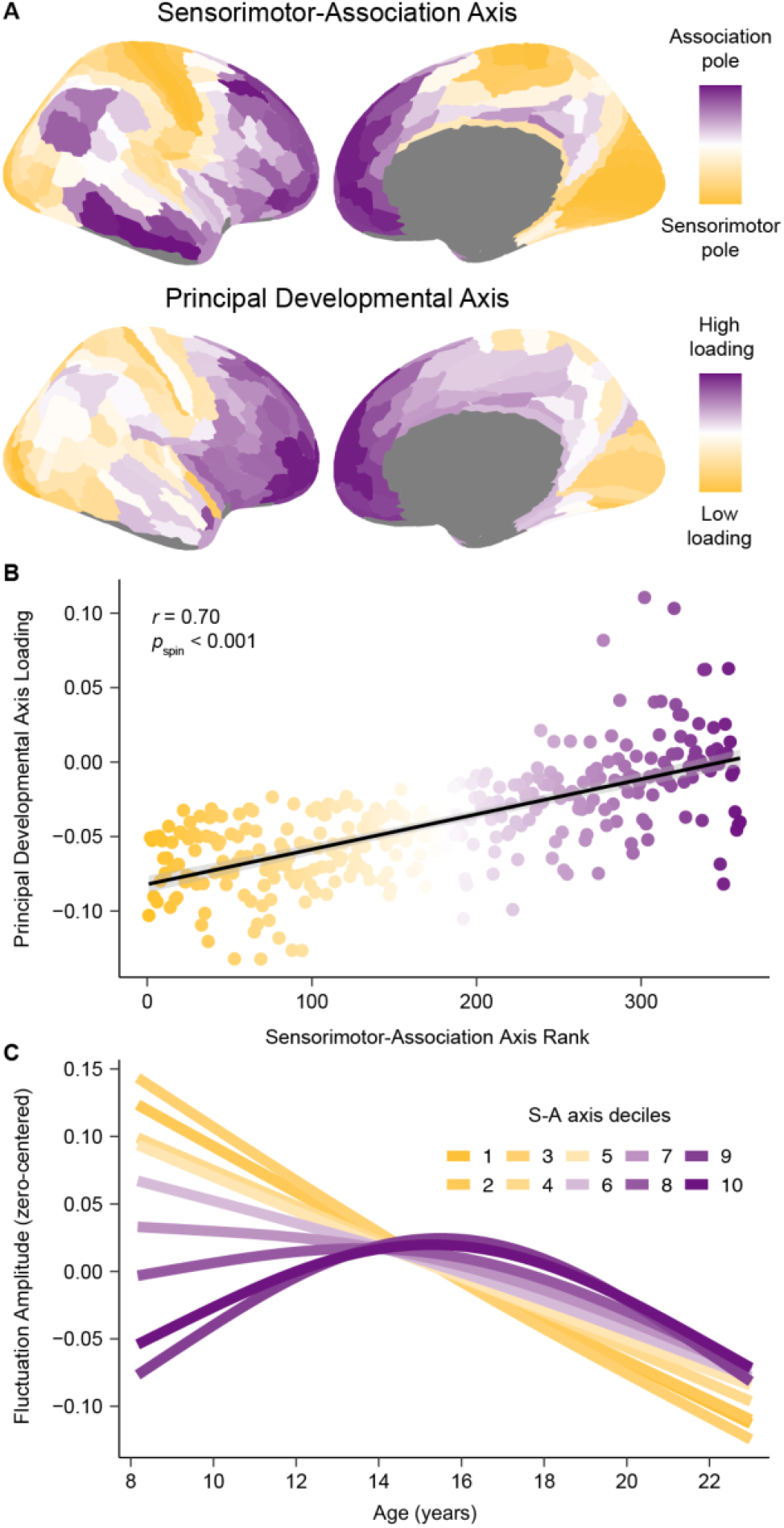
The principal axis of fluctuation amplitude development exhibits convergent embedding with the sensorimotor-association axis. **A)** The principal axis of fluctuation amplitude development closely resembles the sensorimotor-association (S-A) axis, illustrating that the spatiotemporal maturation of intrinsic cortical fMRI activity aligns to the brain’s global cortical hierarchy. The S-A axis, derived in Sydnor et al. (2021)^1^, is a dominant axis of cortical feature organization that spans from primary sensory and motor cortices (sensorimotor pole; dark yellow), to modality-selective and multimodal cortices, and then to transmodal association cortices (association pole; dark purple). The principal developmental axis is the first component from a PCA conducted on regional fluctuation amplitude maturational trajectories. This component quantitatively captures cortex-wide differences in maturational patterns along a unidimensional spatial gradient. **B**) Across the cortex, principal developmental axis loadings strongly correlated with S-A axis ranks. **C**) Average model fits depicting the association between fluctuation amplitude and age are shown for deciles of the S-A axis. To generate average decile fits, the axis was divided into 10 bins each consisting of 33-34 regions, and age smooth functions were averaged across all regions in a bin. The first decile (darkest yellow; linear decline) represents the sensorimotor pole of the axis, the tenth (darkest purple; inverted U) represents the association pole of the axis. Maturational patterns diverged most between S-A axis poles and varied continuously between them.

Our PCA of developmental fits suggests that the spatiotemporal maturation of intrinsic cortical activity conforms to the hierarchical organization of the cortex. In support of this conclusion, we confirmed that principal developmental axis loadings additionally correlated with cortex-wide anatomical (*r* = -0.61), functional (*r* = 0.60), and evolutionary hierarchies (*r* = 0.32). However, the principal developmental axis was significantly more correlated with the S-A axis than with these three hierarchies, which were defined using unimodal data (*p* < 0.001 for all three statistical tests comparing the magnitude of two dependent, overlapping corrections). Neurodevelopmental trajectories were therefore most parsimoniously captured by the S-A axis, which combines information from all three cortical hierarchies and multiple additional data types. To further illustrate the manner in which developmental trajectories for fluctuation amplitude evolve from the sensorimotor to the association end of the S-A axis, we divided the axis into 10 bins and averaged age fits across all regions in a bin. The continuous spectrum of developmental trajectories visible at the regional level (**Fig. 1B**) was recapitulated by S-A axis deciles (**Fig. 3C**), underscoring the extent to which developmental variability was captured by this axis.

### Development is hierarchical through adolescence

The above results highlight that between the ages of 8 and 23 years, age-related changes in fMRI-indexed intrinsic activity are governed by the brain’s S-A axis. We next aimed to elucidate whether this neurodevelopmental pattern was most pronounced during a specific age range, or if it was equally present across the different ages studied. To explore these possibilities, we first calculated each cortical region’s rate of change in fluctuation amplitude at 1 month intervals between 8 and 23 years. Notably, visualizing regional rates of change across the S-A axis (**Fig. 4A**) confirmed that pre-adolescent increases in fluctuation amplitude were uniquely confined to higher-order association cortices. Using these data, we next performed an age resolved analysis where, for each month, we calculated the correlation between regional rates of change and regional S-A axis ranks. This procedure generates age-specific correlations that quantify the extent to which maturational change is spatially ordered along the hierarchy of the S-A axis at a given developmental timepoint. This analysis revealed a robust correlation between developmental change and S-A axis rank from age 8 to 17 years (**Fig. 4B-C**). A maximal correlation value of r = 0.68 (95% credible interval: 0.66 to 0.70) was observed at age 15.0 years (95% credible interval: 14.7 to 15.3 years), indicating peak alignment between neurodevelopment and the S-A axis in mid-adolescence. However, following this peak, the correlation between regional age effects and S-A axis position rapidly declined, dropping to 0 by age 19.3 years (95% credible interval: 18.7 to 20.2 years). These findings reveal that the brain’s developmental program is hierarchical through late adolescence. Following adolescence, however, there may be a programmed switch in the spatial patterning of subsequent age-related change.

**Fig. 4.**
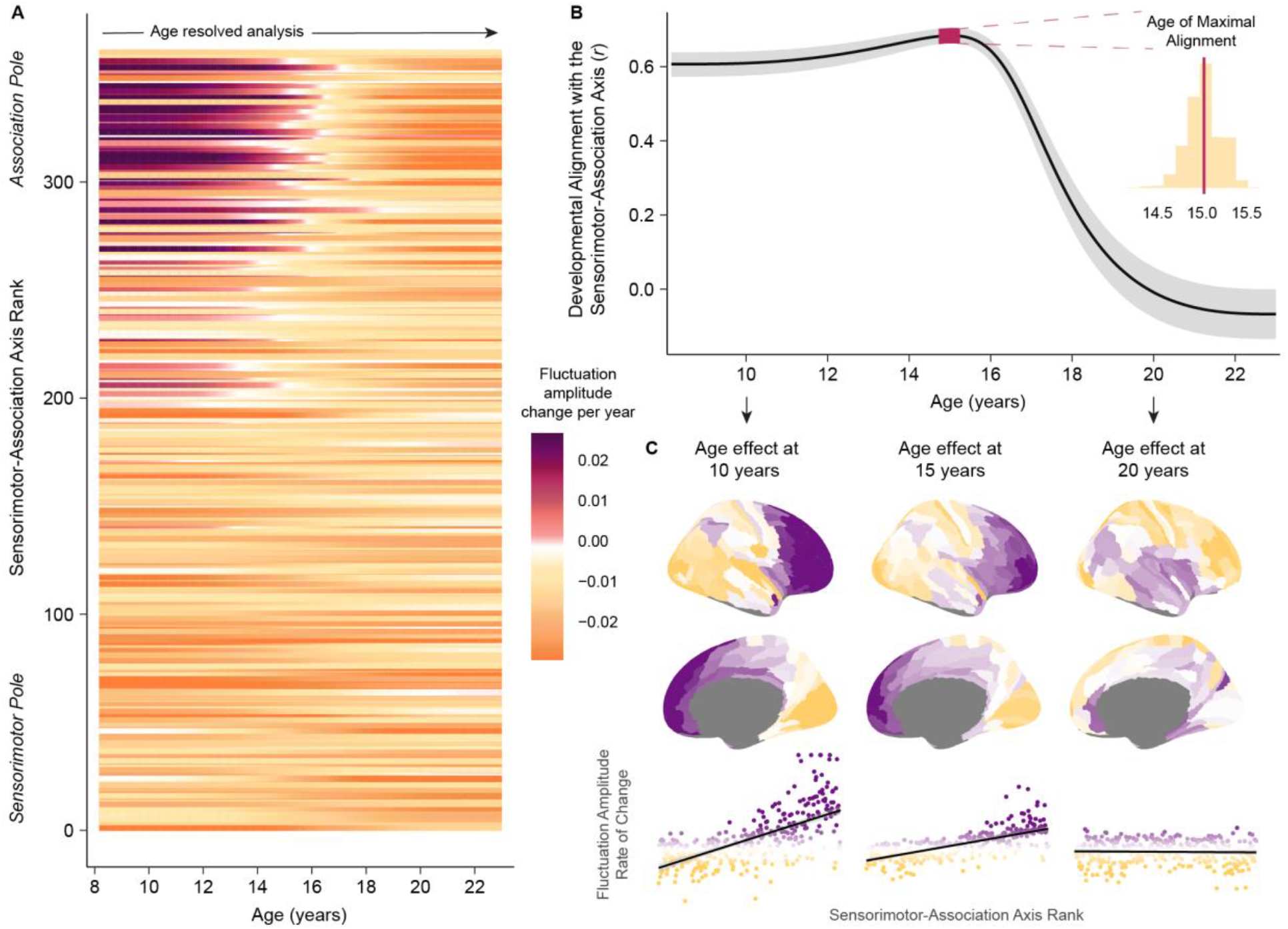
Neurodevelopment progresses along the sensorimotor-association axis until late adolescence. **A)** The rate and direction of developmental change in fluctuation amplitude is displayed from ages 8 to 23 years for each cortical region. Regions are ordered along the y-axis by S-A axis rank. Fluctuation amplitude rate of change, expressed as the change in amplitude per year, was estimated from the first derivative of the GAM smooth function for age. Cortical regions near the association pole of the S-A axis exhibited unique increases in fluctuation amplitude through childhood that culminated in adolescent BOLD amplitude peaks. **B)** Maturational changes in intrinsic BOLD activity align with the S-A axis from childhood until late adolescence. The line plot displays age-specific correlation values (*r*) between regional rates of fluctuation amplitude change and regional S-A axis ranks from ages 8 to 23 years. To obtain reliable estimates of this correlation value at each age, we sampled 10,000 draws from the posterior derivative of each region’s GAM smooth function. We then quantified age-specific correlations between regional derivatives and S-A axis ranks for all 10,000 draws. The median correlation value obtained across all draws is depicted by the black line and the 95% credible interval around this value is represented by the gray band. We additionally determined the age of maximal alignment between fluctuation amplitude change and S-A axis rank for all 10,000 draws. The 95% credible interval for the age of maximal alignment is depicted on the line plot by the pink band, and the full distribution of ages obtained from all draws is portrayed in the inset histogram. **C)** Age-specific developmental effects (first derivative maps) are shown on the cortical surface at age 10 years, 15 years, and 20 years above scatterplots that display the relationship between regional S-A axis ranks (x-axis) and regional age-specific rates of change (y-axis). Scatterplot points are colored by age-specific rates of change. Developmental refinement of fluctuation amplitude was governed by the S-A axis at ages 10 and 15 years. By age 20, further refinement of fluctuation amplitude was unrelated to the S-A axis.

### Developmental results are robust to methodological variation

To ensure that the developmental effects observed were robust to methodological variation and potential confounds, we performed six sensitivity analyses. We evaluated if age-dependent changes in our *in vivo* measure of intrinsic cortical activity were driven by in-scanner head motion, medication use, local cerebral blood flow, regional mean signal intensity, global amplitude differences, or our choice of cortical atlas. In the first two sensitivity analyses, regional GAMs were rerun in the two thirds of the sample with the lowest in-scanner head motion (low motion sample; n = 690; **Fig. 5A**) and in a sample that excluded individuals with current psychoactive medication use or a history of psychiatric hospitalization (no psychiatric treatment; n = 893; **Fig. 5B**). In the next two sensitivity analyses, regional GAMs were refit while additionally controlling for regional cerebral blood flow estimated from arterial spin labeling data (vascular control; **Fig. 5C**) or regional mean T2* signal intensity (T2* signal control; **Fig. 5D**). In the final two sensitivity analyses, regional GAMs were refit with whole brain mean-normalized fluctuation amplitude (mean normalization; **Fig. 5E**) or fluctuation amplitude averaged within Schaefer-400 atlas regions as the dependent variables (atlas replication; **Fig. 5F**).

**Fig. 5.**
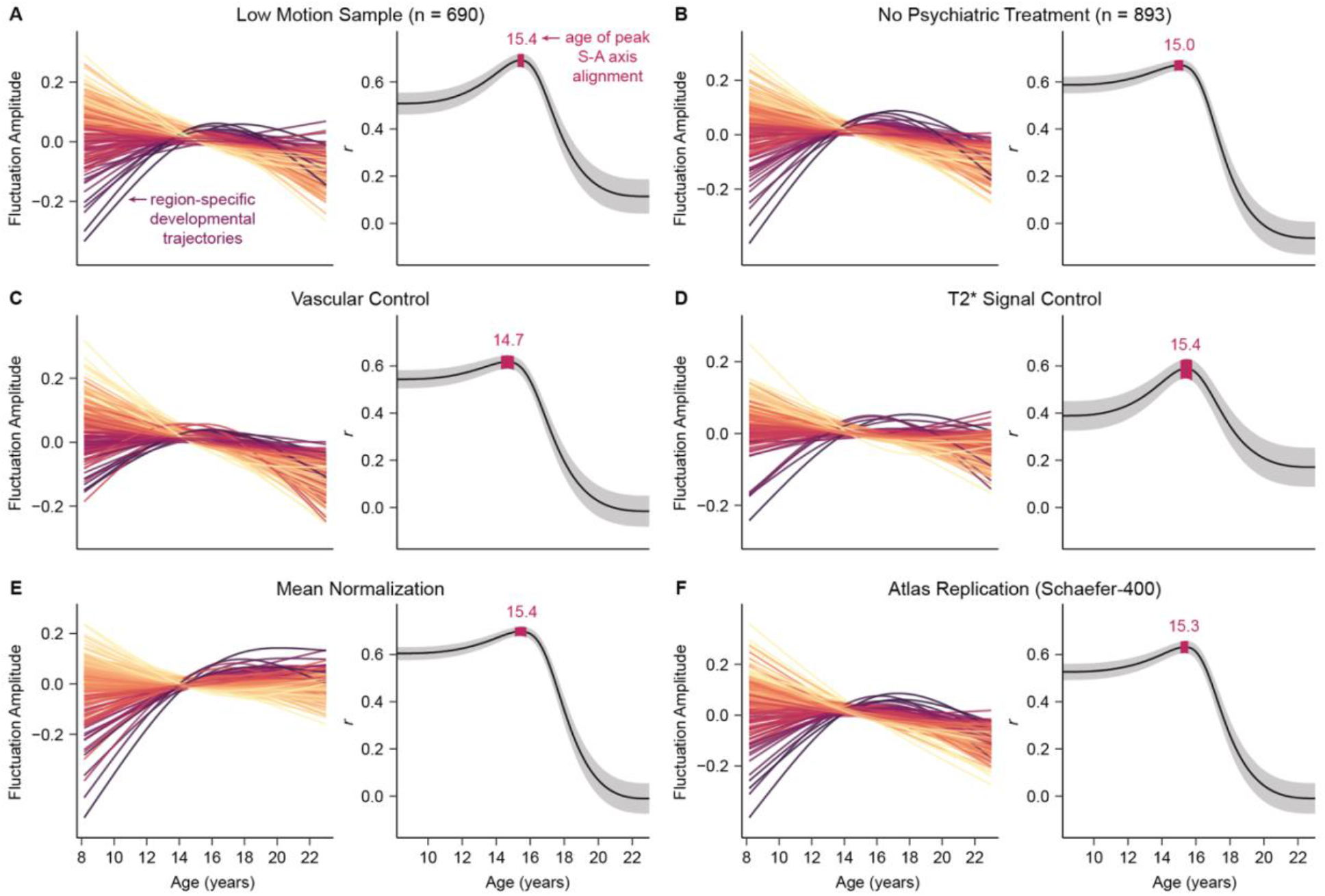
Region-specific and cortex-wide developmental patterns are robust to methodological variation. **A-F)** Key results are shown for each of the six sensitivity analyses performed. For each analysis, the left plot shows fluctuation amplitude developmental trajectories (zero-centered) for left hemisphere regions, colored by age effects. The right plot presents the age-dependent analysis of the association between developmental change in fluctuation amplitude and S-A axis rank. All six sensitivity analyses yielded convergent region-specific and cortex-wide results, confirming that our developmental findings were not being driven by head motion in the scanner (A), the use of psychotropic medications (B), age-related changes in cerebrovascular perfusion (C), inter-scan differences in T2* signal strength (D), global effects (E), or the specific atlas used for cortical parcellation (F).

In each of the six sensitivity analyses, region-specific fluctuation amplitude maturational trajectories very closely mirrored the developmental fits from the main analysis, with negative age effects observed in most cortices but positive effects seen in select transmodal association cortices. Consequently, regardless of the sample used or controls performed, a cortical region’s age effect was fundamentally and significantly (all *p*_*s*pin_ < 0.05) related to its position in the S-A axis (main analysis: *r* = 0.54, low motion sample: *r* = 0.56, no psychiatric treatment: *r* = 0.51, vascular control: *r* = 0.51, T2* signal control: *r* = 0.49, mean normalization: *r* = 0.66, atlas replication: *r* = 0.43). Furthermore, for all sensitivity analyses, our age resolved analysis confirmed a strong correlation between the rate of fluctuation amplitude change and S-A axis rank from childhood to late adolescence, with the peak age of neurodevelopmental alignment to this axis occurring during early adolescence. These analyses verify that our findings concerning the nature and patterning of age-dependent changes in cortical fMRI activity are robust to methodological variation.

### Environmental influences on intrinsic fMRI fluctuations diverge across the sensorimotor-association axis

Instrumental work in animal models has shown that environmental exposures can affect the development of plasticity-regulating mechanisms, including the maturation of inhibitory interneurons^52^ and intracortical myelination^53,54^. Given these preclinical findings, we sought to investigate the hypothesis that in youth, environmental influences become embedded in the brain by affecting cortical plasticity. We tested this theory by examining relationships between inter-individual variability in children’s environments and individual differences in our in vivo measure of activity-indexed plasticity. Multivariate features of each participant’s neighborhood environment were summarized using a single previously-published factor score^55^. Higher factor scores indicate that an individual lived in a neighborhood with a higher median family income, lower population density, fewer vacant housing lots, a greater percentage of residents who are married, employed, and high school educated, and a lower percentage of residents in poverty (**Fig. 6A**). Associations between environment factor scores and regional fluctuation amplitude were modeled with GAMs while controlling for developmental effects and other covariates (in-scanner motion and sex).

**Fig. 6.**
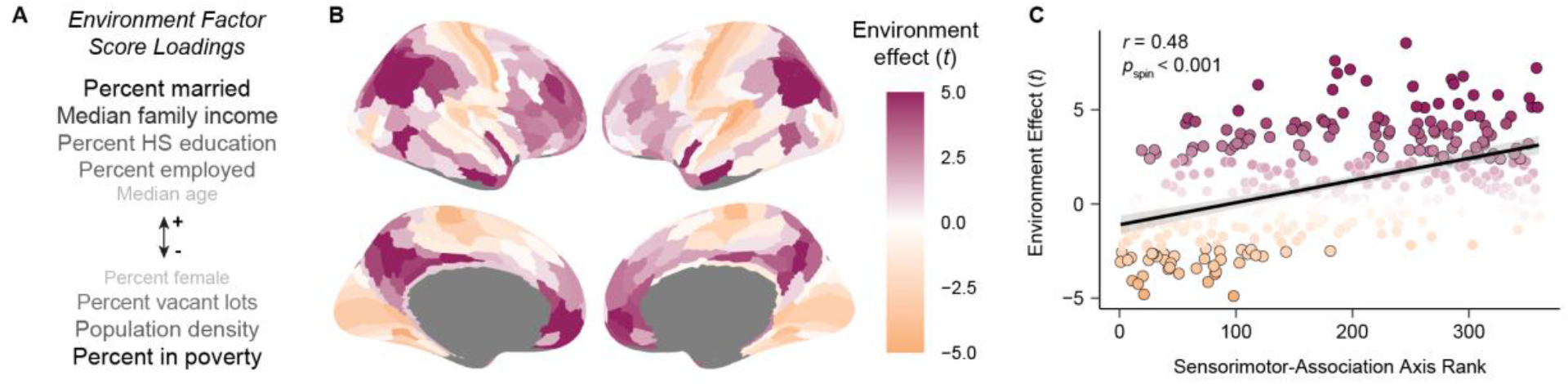
Associations between fluctuation amplitude and the developmental environment vary along the sensorimotor-association axis. **A**) The environment factor score captures multiple features of the neighborhood environment each child lived in. Variables listed above the arrow (+) positively loaded onto the environment factor score. Variables listed below the arrow (-) negatively loaded onto the factor score. Darker and larger text indicates stronger loadings. **B**) The effects of the environment on regional fluctuation amplitude. The cortical map displays the main effect of environment factor score (*t*-values) from a GAM that included age, sex, and in-scanner motion as covariates. Positive values (magenta) designate positive associations whereas negative values (orange) indicate negative associations. Lower environment factor scores—representing higher poverty, higher population density, and lower high school (HS) education and employment rates at the neighborhood level—were associated with lower fluctuation amplitude in association cortex but higher fluctuation amplitude in primary sensorimotor cortices. **C**) The S-A axis explains significant variability in the effects of the environment on intrinsic fMRI activity. Data points are colored by environment effect *t*-values; black outlines denote statistically significant effects.

Forty-two percent of cortical regions showed a significant association between fluctuation amplitude and youth’s neighborhood environments (*p*_FDR_ < 0.05 in 141 regions). Higher environment factor scores were associated with higher fluctuation amplitude across the association cortex, but with lower fluctuation amplitude nearly exclusively within primary and early sensory and motor cortices (**Fig. 6B**). This pattern of effects indicates that youth raised in neighborhoods with higher income, education, and employment rates and with lower population density and poverty tended to have greater amplitude intrinsic fMRI fluctuations in higher-order cortices and lower amplitude fluctuations in modality-specific cortices. This divergence in environment effects intimated that relationships between the developmental environment and cortical intrinsic activity could systematically vary along the brain’s S-A axis. Supporting this notion, environment effects and S-A axis rank were significantly correlated (*r* = 0.48, *p*_spin_ < 0.001; **Fig. 6C**). Overall, our analysis of youth’s neighborhoods suggests that the developmental environment may impact cortical intrinsic activity, with the nature of the impact differing across a principal axis of brain organization and development.

## DISCUSSION

During embryonic and early postnatal cortical development, developmental programs are spatially and temporally governed by major organizing axes. Cortical arealization is controlled by transcription factors expressed along anterior-medial to posterior-lateral axes^56–58^ and neurogenesis terminates along an anterior-posterior axis^59^. The alignment of developmental programs with neuroaxes is thus a fundamental element of early cortical development. In the current study, we demonstrate that the maturation of intrinsic cortical activity conforms to the hierarchical S-A axis from ages 8 to 18 years, thus establishing that this core facet of development extends to childhood and adolescence. Specifically, we demonstrate that declines in the amplitude of intrinsic fMRI activity occur during childhood and adolescence, are temporally coupled to the increasing expression of a plasticity-restricting factor, and are temporally staggered along the S-A axis of cortical organization. We additionally show that the S-A axis captures not only inter-regional variation in cortical maturational profiles, but also variation in the effects of children’s developmental environments on intrinsic activity dynamics. Together, these results provide evidence of a hierarchical axis of neurodevelopmental plasticity in youth.

Intrinsic activity has a profound influence on the immature brain, impacting neuron survival, circuit wiring, topographic map formation, synaptic connectivity, and overall cortical volume^22,60–64^. Prominent changes in the prevalence and patterning of this activity occur during development, engendered by shifts in the maturity of the cortex^23–25^ and in the level of cortical plasticity^15,21,26^. Here, we aimed to characterize developmental changes in intrinsic activity with resting-state fMRI and observed prolonged declines in the amplitude of BOLD activity— potentially reflecting decreases in spontaneous activity-indexed plasticity—throughout the protracted course of human neurodevelopment. Notably, declines in activity amplitude occurred heterochronously across the cortical mantle, highlighting temporal developmental variability. In general, unimodal sensory and motor regions exhibited large and early declines in fluctuation amplitude, which began around 8 to 12 years of age. Select association cortices underwent later-onset amplitude declines beginning typically between 13 to 16 years. In contrast, only transmodal association cortices displayed periods of increasing activity amplitude; increases lasted until adolescence and were followed by significant declines. These region-specific windows of declining fluctuation amplitude occur after peak gray matter volume is obtained^3^ and coincide with regional windows of extensive synaptic pruning^7,65^. In addition, we demonstrated that the onset of decline in fluctuation amplitude in each region was linked to its period of maximal intracortical myelin growth. Maturational refinement of this non-invasive measure of intrinsic cortical activity thus appears to be temporally coupled to developmental changes in multiple markers of circuit plasticity, suggesting the presence of a gradient of neurodevelopmental plasticity during childhood and adolescence.

Indeed, in support of our proposed theory on developmental chronology^1^, we found that regional differences in the development of fluctuation amplitude were systematically explained by the asynchronous patterning of cortical change across the S-A axis. Regional developmental trajectories diverged in a continuous and graded manner across this axis, with reductions in the amplitude and synchrony of intrinsic activity appearing to occur earliest and most extensively in regions ranked lowest in the axis. This pattern of results reveals that regional maturational profiles are organized by the brain’s global cortical hierarchy, and furthermore suggest that cortically plasticity may progressively decline along this hierarchy with age. Our data also reveal that a functional signature observed during early periods of plasticity increases in transmodal association cortices until the middle of adolescence, indicating there may be a heightened period of developmental plasticity in these cortices during late development. This finding reinforces theories that adolescence is a sensitive or critical period for the development of association cortex during which the environment, experience, and interventions could have a large impact on malleable higher-order cortices^1,14,66^.

Although non-invasive imaging data cannot establish the mechanisms underlying this S-A developmental pattern, prior work suggests that it may emerge due to the hierarchical maturation of key plasticity-regulating neurobiological events. Developmental plasticity is regulated by a host of neurochemical and structural mechanisms^16^. Key mechanisms include the local strength of cortical inhibition^28,67^, the patterning of thalamocortical input^22,68^, and the density of both intracortical myelin^30^ and perineuronal nets^21,69^. Critically, these mechanisms mature earlier in sensory cortices and later in association cortices^14,15,70–72^. Moreover, these mechanisms elicit transitions in the patterning of intrinsic cortical activity, impacting the duration, strength, or synchronicity of activity^21,27,29,49,73,74^. Hierarchical maturation of plasticity-regulating features may therefore be responsible for waves of changing intrinsic activity dynamics along the S-A axis. Given that alignment of developmental change with the S-A axis was maximal in mid-adolescence and continued until only 18 years of age, these developmental plasticity mechanisms may account for changes in intrinsic activity during the first two decades of life, but not after.

Animal studies of cortical plasticity have established that enriched versus deprived developmental environments can differentially affect the maturation of the aforementioned plasticity-regulating features, including the maturation of inhibitory interneurons^52,75^, intracortical myelination^53,54^, and perineural nets^75^. Data from human studies additionally suggest that the level of unpredictability, stress, and adversity in children’s environments influences the pace of brain development^42–44^, providing convergent support for an effect of the environment on the expression of cortical plasticity in youth. The present study complements past work by showing that variability in children’s neighborhood environments is associated with differences in a proposed functional correlate of ongoing circuit plasticity. Strikingly, the magnitude and direction of brain-environment associations observed here were stratified by S-A axis rank, signifying that the environment may exert differential effects on cortical regions dependent upon each region’s developmental trajectory. In higher-order association cortices, youth that lived in neighborhoods with greater poverty, unemployment rates, and population density had a lower amplitude of intrinsic fMRI fluctuations. This directional effect is potentially suggestive of a reduced potential for plasticity in late-maturing cortices, and thus coheres with leading theories that environmental adversity hastens the pace of cortical maturation^42,44^. Yet, youth from neighborhoods with lower social capital and community resources additionally exhibited higher fluctuation amplitude in early-maturing sensorimotor cortices—consistent with a more immature cortex. We speculate that this series of results could reflect a hastening of developmental timing up the hierarchical S-A axis in youth from disadvantaged neighborhoods, putatively resulting in higher expression of plasticity-limiting features early in association cortices at the expense of limiting plasticity in primary regions.

Several limitations of this work should be noted. First, this was a cross-sectional investigation of neurodevelopment in youth. Future investigations with longitudinal study designs could characterize within-individual changes in cortical intrinsic activity as well as the effects of the environment on the pace of an individual’s development. Longitudinal studies will also be better suited to examine temporal precedence between developmental refinement of intrinsic fMRI activity and maturation of plasticity-regulating features. Second, we used resting-state functional MRI to study intrinsic cortical activity, however the BOLD signal is sensitive to neural, vascular, and respiratory factors. While sensitivity analyses aimed at addressing these factors provided highly convergent results, future studies using more direct measures of neural activity (e.g., electrocorticography) may therefore be helpful for extending the present results. Third, this work focused on the cortex only. Assessing whether neurodevelopment proceeds hierarchically from subcortical areas supporting sensory and motor functions to those supporting cognitive and socioemotional functions is a key avenue for future exploration.

The present study demonstrates that during childhood and adolescence, the spatiotemporal patterning of development in intrinsic cortical fMRI activity coheres with a hierarchical axis of cortical organization. This result emphasizes that the S-A axis is both a dominant spatial feature axis and a core neurodevelopmental axis, and intimates that spatial form in the adult brain may emerge from coordinated development during youth^1,76,77^. In addition, the observed refinements in fMRI fluctuations suggest that shifts in circuit plasticity temporally progress along this axis, calling for studies that can mechanistically establish the neurobiological events driving S-A developmental plasticity^14^. Such events may include the maturation of neurochemical and structural plasticity-regulating features as well as successive expression of molecules that orchestrate developmental timing, for example circadian clock genes or other temporally organized transcription factors^15,78,79^. Given the relevance of the S-A axis for understanding cortical development in childhood and adolescence, future work should explore whether major organizing axes play a role in cortical refinement during infancy and early childhood. Continued discovery of temporal axes of development across human’s multi-decade maturational course will provide evidence of how plasticity is distributed across brain regions at different developmental stages. Such insights into the temporal patterning of plasticity may help to guide interventions in youth that align with each child’s neurotemporal context.

## METHODS

### Participants

Participants were recruited as part of the Philadelphia Neurodevelopmental Cohort^80^, a community study of child and adolescent brain development. Demographic, clinical, environmental, and neuroimaging data from 1033 youth were included in the present study.

Study sample demographics include an age range of 8 to 23 years (mean age = 15.7 ± 3.3 years), a sex distribution of 467 males and 566 females, and a race distribution that was 47% White, 41% Black, 11% identifying with more than one race, 7% Asian, and 0.3% US Indian or Alaskan Native. All participants over the age of 18 gave written informed consent prior to study participation. Participants under the age of 18 gave informed assent with written parental consent. All study procedures were approved by the Institutional Review Boards of the University of Pennsylvania and the Children’s Hospital of Philadelphia.

### MRI data acquisition

T1-weighted structural MRI and resting-state functional MRI data were used in the present study. All MRI scans were acquired on the same 3T Siemens Tim Trio scanner at the University of Pennsylvania with a 32-channel head coil. T1-weighted structural images were acquired with a magnetization-prepared rapid acquisition gradient-echo (MPRAGE) sequence with the following parameters: repetition time = 1810 ms, echo time = 3.51 ms, inversion time = 1100 ms, flip angle = 9 degrees, field of view = 180 × 240 mm, matrix = 192 × 256, slice number = 160, voxel resolution = 0.94 × 0.94 × 1 mm). Resting-state functional images were acquired with a single-shot, interleaved multi-slice gradient-echo echo planar imaging (GE-EPI) sequence with the following parameters: repetition time = 3 s, echo time = 32 ms, flip angle = 90 degrees, field of view = 192 × 192 mm, matrix = 64 × 64, slice number = 46, voxel resolution = 3 mm^3^, volumes = 124. To enable susceptibility distortion correction of resting-state functional images, a map of the main magnetic field (i.e., a B0 field map) was additionally collected using a dual-echo, gradient-recalled echo (GRE) sequence with the following parameters: repetition time = 1000 ms, echo time 1 = 2.69 ms, echo time 2 = 5.27 ms, flip angle = 60 degrees, field of view = 240 × 240 mm, matrix = 64 × 64, slice number = 44, voxel resolution = 3.8 × 3.8 × 4 mm.

### MRI data processing

T1-weighted images and resting-state functional MRI timeseries were processed with fMRIPrep 20.2.3^81^. The T1-weighted image was corrected for intensity non-uniformity with Advanced Normalization Tools’ (ANTs 2.3.3) N4BiasFieldCorrection^82^, skull stripped with a Nipype implementation of the ANTs brain extraction workflow, tissue segmented with FSL fast 5.0.9, and used for cortical surface reconstruction with FreeSurfer 6.0.. The T1-weighted image was additionally non-linearly registered to the MNI152 T1 template (volume-based spatial normalization) with ANTs.

To preprocess functional scans, a skull-stripped reference BOLD volume was first generated and a B0 fieldmap was co-registered to this reference volume. The B0 field map was estimated based on the phase-difference map calculated with the dual-echo GRE sequence, converted to a displacements field map with FSL’s fugue and SDCflow tools, and used for susceptibility distortion correction of the reference BOLD volume. The susceptibility corrected BOLD reference was then rigidly co-registered (6 degrees of freedom) to the T1 reference using boundary-based registration implemented with FreeSurfer’s bbregister^83^. The functional MRI timeseries were slice-time corrected using 3dTshift from AFNI 20160207 and then resampled onto their original, native space by applying a single, composite transform to correct for susceptibility distortions and for in-scanner head motion. Head motion parameters were calculated with respect to the reference BOLD volume prior to any spatiotemporal filtering using FSL mcflirt; six rotation and translation parameters were calculated. BOLD timeseries were additionally resampled into standard space, generating preprocessed timeseries in the MNI152 T1 template, and onto the fsaverage surface. Finally, to project functional timeseries onto the fsLR cortical surface for study analyses, grayordinates files containing 32k vertices per hemisphere were generated using the highest-resolution fsaverage as an intermediate standardized surface space. Volumetric resampling was performed using antsApplyTransforms, configured with Lanczos interpolation to minimize the smoothing effects of other kernels. Surface resampling was performed using FreeSurfer mri_vol2surf.

fMRIPrep was additionally used to estimate the following 36 confounds from the preprocessed timeseries: six head motion parameters; three region-wise global signals (mean cerebrospinal fluid, white matter, and whole brain signals); temporal derivatives of the six head motion parameters and the three global signal estimates; and quadratic terms for the motion parameters, tissue signals, and their temporal derivatives^84–86^. These confound matrices were utilized within xcp_abcd 0.0.4, which is an extension of the top-performing eXtensible Connectivity Pipeline (XCP) Engine^84,85^ specifically developed to mitigate motion-related artifacts and noise in resting-state functional MRI data from developmental samples. With xcp_abcd, preprocessed functional timeseries on the fsLR cortical surface underwent nuisance regression using the 36 confounds listed above. Confounds were regressed using linear regression as implemented in Scikit-Learn 0.24.2.

### Fluctuation amplitude quantification

To measure the relative level and synchronization of spontaneous fMRI activity, we quantified fluctuation amplitude, defined as the power of low frequency BOLD fluctuations. To calculate fluctuation amplitude, processed fsLR surface BOLD timeseries were first transformed from the time domain to the frequency domain and a power spectrum was generated in the 0.01-

0.08 Hz range. The mean square root of the power spectrum was then calculated; the mean square root represents the average amplitude (intensity) of time-varying resting-state BOLD fluctuations within this low frequency band^39^. Simultaneous BOLD and electrophysiological or calcium recordings have revealed that slow fluctuations in BOLD signal are coupled to fluctuations in intra-cortically measured neural signals, including both local field potentials and infra-slow calcium signals^33–38^. Of note, the fluctuation amplitude measure used here is analogous to other commonly used spectral- or variability-based BOLD measures, including the amplitude of low frequency fluctuations (ALFF) and resting-state functional amplitude (RSFA).

Fluctuation amplitude was quantified at the vertex-level and then parcellated with fsLR surface atlases to provide mean fluctuation amplitude within individual cortical regions. The HCP multimodal atlas^87^ was used for all primary analyses and the Schaefer-400 atlas^88^ was used for a sensitivity analysis. Parcellation was conducted in R with the ciftiTools package^89^ utilizing Connectome Workbench 1.5.0^90^. Fluctuation amplitude was not analyzed within cortical regions that exhibited low signal to noise ratio (SNR) in >= 25% of their assigned vertices. Low SNR vertices were defined identically to our prior work^91^ as vertices with an average (across-participant) BOLD signal < 670 after normalizing signal to a mode of 1000. Twenty-four parcels located within the orbitofrontal cortex and the ventral temporal lobe were excluded from both the HCP multimodal atlas and the Schaefer-400 atlas.

### MRI sample construction

1374 individuals in the Philadelphia Neurodevelopmental Cohort had T1-weighted images, B0 field maps, and identical parameter^92^ resting-state functional MRI scans available. From this original sample of n = 1374, 120 individuals were excluded from the study due to medical problems that could impact brain function or incidentally encountered abnormalities of brain structure^93,94^. Data from 202 additional participants were excluded due to low quality T1-weighted images and FreeSurfer reconstructions (n = 23) or high in-scanner head motion (n = 179). As in prior work, high in-scanner head motion was defined as a mean relative root mean squared framewise displacement > 0.2 mm during the functional scan (Satterthwaite et al., 2012). Using data from the remaining sample (n = 1052), we identified fluctuation amplitude outliers at the regional level based on a cut off of ± 4 standard deviations from the mean. Individuals with outlier data in more than 5% of cortical regions (n = 19) were excluded, producing the final study sample of 1033 individuals.

### Characterizing developmental effects

#### Generalized additive models

All statistics were carried out in R 4.0.2. In order to flexibly model linear and non-linear relationships between fluctuation amplitude and age, we implemented GAMs using the mgcv package in R^95^. GAMs were fit with regional fluctuation amplitude as the dependent variable, age as a smooth term, and sex and in-scanner motion as linear covariates. Models were fit separately for each parcellated cortical region using thin plate regression splines as the smooth term basis set and the restricted maximal likelihood approach for smoothing parameter selection. The GAM smooth term for age produces a spline, or a smooth function generated from a linear combination of weighted basis functions, that represents a region’s developmental trajectory. To prevent overfitting of the spline, we set the maximum basis complexity (k) to 3 to limit the number of basis functions that could be used to estimate the overall model fit. A value of k = 3 was chosen over higher values (i.e., k = 4-6) given that this basis complexity resulted in the lowest model Akaike information criterion (AIC) for the vast majority of cortical regions. Statistical tests of the k-index^95^, which estimate the degree of unaccounted for pattern in the residuals, confirmed that this basis dimension was sufficient.

For each regional GAM, the significance of the association between fluctuation amplitude and age was assessed through an analysis of variance (ANOVA) that compared the full GAM model to a nested, reduced model with no age term. A significant result indicates that the residual deviance was significantly lower when a smooth term for age was included in the model, as assessed with the chi-square test statistic. We corrected ANOVA *p*-values across all region-wise GAMs using the false discovery rate (FDR) correction and set statistical significance at *p*_FDR_ < 0.05. For each regional GAM with a significant age smooth term, we furthermore identified the specific age range(s) wherein fluctuation amplitude was significantly changing through the *gratia* package in R. Age windows of significant change were identified by quantifying the first derivative of the age smooth function (Δ fluctuation amplitude / Δ age) using finite differences and determining when the simultaneous 95% confidence interval of this derivative did not include 0^96^. To establish the overall magnitude and direction of the association between fluctuation amplitude and age, which we refer to throughout as a region’s overall age effect, we calculated the partial *R*^*2*^ between the full GAM model and the reduced model (effect magnitude) and signed the partial *R*^*2*^ by the sign of the average first derivative of the smooth function (effect direction). Finally, for each cortical region, we also explored whether fluctuation amplitude developmental trajectories significantly differed between males and females by adding a factor-smooth interaction term to the main GAM model. Following FDR correction for multiple comparisons across all region-wise GAMs, no significant interaction terms were observed; sex effects were therefore not evaluated further.

#### Associations with cortical myelin development

We formally assessed whether the development of intrinsic fMRI activity amplitude was spatially and temporally related to the development of intracortical myelination. Cortical myelin development was previously comprehensively characterized by Baum et al. (2022)^70^ using T1w/T2w surface-based myelin mapping^46,97^ in a sample of 628 youth ages 8 to 21 years who had data collected as part of the Human Connectome Project in Development. Using high resolution (0.8 mm^3^) T1-weighted and T2-weighted images, HCP processing pipelines, and state of the art methods for B1+ transmit field bias correction, partial volume reduction, and CSF correction^97^, Baum et al. (2022)^70^ investigated the maturational trajectory of increases in the T1w/T2w ratio in each region. In this investigation, the authors fit region-specific GAMs with a smooth term for age using thin plate regression splines as the smoothing basis, paralleling the present work. GAMs included covariates for sex, scanner, and B1+ transmit field correction-related variables, following current best practices^97^ for statistically comparing the T1w/T2w ratio across individuals.

To test if the extent to which fluctuation amplitude changed with age was related to the degree to which cortical myelin content changed with age, we calculated the correlation coefficient between the two distinct age effects: those calculated from regional fluctuation amplitude GAMs and those calculated from regional T1w/T2w ratio GAMs reported by Baum et al. (2022)^70^. As in the present work, T1w/T2w ratio age effects were determined by a partial *R*^*2*^ derived by comparing the full GAM model to a reduced model with no smooth term for age. The association between fluctuation amplitude age effects and T1w/T2w ratio age effects was quantified with a Spearman’s correlation coefficient. A Spearman’s correlation was also used to assess whether there was temporal correspondence between the development of fluctuation amplitude and the T1w/T2w ratio across the cortex. Specifically, we calculated the correlation coefficient between the age at which fluctuation amplitude began to significantly decrease in each region and the age at which the T1w/T2w ratio maximally increased. The age at which fluctuation amplitude began to significantly decrease was the first age at which the derivative of the age smooth function was significantly negative. The age at which the T1w/T1w ratio maximally increased was the age at which the derivative of the age smooth function was maximal.

#### Alignment with the sensorimotor-association axis

This work set out to test the overarching hypothesis that neurodevelopmental patterns are related to the S-A axis during childhood and adolescence. We therefore examined whether patterns of fluctuation amplitude maturation aligned with the S-A axis developed in our prior work^1^. The S-A axis was derived by averaging rank orderings of ten cortical feature maps that exhibit systematic variation between lower-order sensorimotor cortices, middle-order unimodal and multimodal cortices, and higher-order heteromodal and paralimbic association cortices^1^. These maps include the functional hierarchy delineated by the principal gradient of functional connectivity^20^, the evolutionary hierarchy defined by macaque-to-human cortical areal expansion^51^, the anatomical hierarchy as quantified by the T1w/T2w ratio^50^, allometric scaling calculated as local areal scaling with scaling of total brain size^98^, aerobic glycolysis measures of brain metabolism^99^, cerebral blood flow measures of brain perfusion^100^, gene expression patterning indexed by the principal component of brain expressed genes^50^, a primary mode of brain function characterized by the principal component of NeuroSynth meta-analytic decodings^101^, a histological gradient of cytoarchitectural similarity developed using the BigBrain atlas^102^, and cortical thickness measured by structural MRI. The resulting S-A axis represents a dominant, large-scale motif of cortical organization that captures the stereotyped patterning of cortical heterogeneity from primary visual, auditory, and somatosensory regions (lowest ranks in the S-A axis) to transmodal frontal, temporal, and parietal association regions (highest ranks in the S-A axis).

We performed the following analyses to ascertain whether the development of fluctuation amplitude may indeed be governed by the hierarchical S-A axis. Using Spearman’s correlations, we evaluated associations between cortical regions’ S-A axis ranks and both 1) their magnitude of fluctuation amplitude development (GAM age effect) and 2) the age at which their fluctuation amplitude began significantly decreasing (first significant negative derivative). We next conducted a PCA on regional developmental trajectories. The goal of this PCA was to visualize the spatial axis that explained the greatest variance in how an *in vivo* measure of cortical intrinsic activity changed with age. The input to the PCA was region-wise age fits (zero-averaged smooth function estimates). The first principal component generated by this PCA contained regional loadings that capture differences in maturational patterns across one low-dimension embedding. We quantified the similarity between the first principal component (loadings) and the cortex’s S-A axis^1^, anatomical hierarchy^50^, functional hierarchy^20^, and evolutionary hierarchy^51^ with independent Spearman’s correlations. We additionally assessed whether the correlation with the S-A axis was significantly greater in magnitude than correlations with the three hierarchies using a statistical test for comparing two dependent, overlapping correlations that utilizes a back-transformed average Fisher’s Z procedure ^103,104^.

Finally, we implemented an age resolved analysis to evaluate if the development of fluctuation amplitude aligned with the S-A axis throughout the entire developmental window studied. For this analysis, we computed across-region Spearman’s correlations between S-A axis rank and the first derivative of the GAM age spline at 200 ages between 8 and 23 years. Hence, we quantified the relationship between a region’s fluctuation amplitude change per year and its position in the S-A axis at 200 age increments—allowing us to study changes in the extent of S-A axis alignment over the course of development. We determined a correlation coefficient point estimate as well as a 95% credible interval for these age-specific correlation values. To do so, we sampled from the posterior distribution of each region’s fitted GAM 10,000 times, generating 10,000 simulated age smooth functions and corresponding derivatives. We then repeated the process of correlating S-A axis rank with the first derivative of the age smooth function at each of the 200 ages for all 10,000 posterior draws, generating a sampling distribution of possible correlation values at each age increment. This distribution was used to calculate the median correlation value and the 95% credible interval of correlation values at each age. In addition, the sampling distribution of age-specific S-A axis correlation values was used to identify the age at which fluctuation amplitude development maximally aligned with the S-A axis and the youngest age at which no alignment to the axis was observed. To discover ages of maximal and null alignment, we calculated the age at which the axis correlation was largest as well as the first age at which the correlation equaled 0 for all 10,000 draws. For both measures, the median age across all draws and a 95% credible interval was calculated.

#### Sensitivity analyses

We performed a series of sensitivity analyses to confirm that the developmental effects observed were not being driven by potentially confounding factors including in-scanner head motion, psychiatric medication use, cerebrovascular perfusion, BOLD signal intensity, global amplitude effects, or the atlas used for cortical parcellation. For each sensitivity analysis, regional GAMs were refit either in a reduced sample (head motion and psychiatric medication analyses), in the full sample but with an additional model covariate (vascular and BOLD signal intensity analyses), or in the full sample with a modified dependent variable (global amplitude normalization and cortical atlas analyses). GAM-derived fluctuation amplitude trajectories were then visualized and developmental alignment with the S-A axis was confirmed.

The first sensitivity analysis was conducted with a low motion sample to mitigate the potential confounding effect of in-scanner head motion on fluctuation amplitude^86^. From the main study sample of 1033 individuals, we excluded 343 individuals with a mean relative root mean squared framewise displacement > 0.075, retaining a low motion sample of n = 690 (ages 8-23 years; mean age = 16.1 years; 395 female). The second sensitivity analysis was carried out to ensure that psychotropic medication use, which was more frequent among older study participants, did not explain the age-related changes in fluctuation amplitude. GAMs were refit after removing all participants (n = 140) from the original sample of 1033 individuals that reported current psychoactive medication use or a history of psychiatric hospitalization (remaining n = 893; ages 8-23 years; mean age = 15.6 years; 507 female).

The third sensitivity analysis aimed to address the fact that the hemodynamic BOLD signal has both neuronal and vascular contributors. Prior work has demonstrated that measures of BOLD fluctuation amplitude contain substantial physiological information not attributable to vascular properties such cerebrovascular reactivity, rigidity, and blood flow^105^. Nonetheless, we still evaluated whether changes in vascular reactivity or cerebral perfusion with age could potentially be contributing to our developmental findings concerning fluctuation amplitude. We approached this evaluation by directly controlling for each participant’s regional cerebral blood flow, a measure of local blood perfusion, in region-wise GAMs. Cerebral blood flow was estimated from arterial spin labeling (ASL) data collected from participants with a pseudo-continuous ASL (pCASL) sequence with the following acquisition parameters: repetition time = 4000 ms, echo time = 2.9 ms, voxel resolution = 2.29 × 2.29 × 6 mm, label duration = 1500 ms, post label delay = 1250 ms, 40 paired label and control acquisition volumes. Data were processed using ASLPrep version 0.2.6 as in Adebimpe et al. (2022)^106^. Standard cerebral blood flow maps were generated and parcellated with the HCP multimodal atlas. Thirty-one participants included in the main study sample did not have ASL data available thus this vascular control analysis was performed using data from the remaining 1002 participants.

The fourth sensitivity analysis was undertaken to rule out the possibility that inter-individual differences in regional mean BOLD signal intensity, rather than BOLD fluctuations per se, could account for our findings. In the full study sample (n = 1033), region-specific GAMs were refit while adding regional mean BOLD signal (i.e., the average T2* signal from minimally preprocessed functional timeseries) as an additional control covariate. Regional mean BOLD signal was calculated from parcellated fsLR surface BOLD timeseries generated with fMRIPrep by averaging the BOLD signal intensity in each parcellated cortical region across all volumes; this measure was calculated prior to regression of confounding signals.

The fifth sensitivity analysis used mean normalized fluctuation amplitude as the dependent variable in all regional GAMs to examine the extent to which region-specific changes in fluctuation amplitude with age occurred above and beyond changes in global mean fluctuation amplitude. This sensitivity analysis was motivated by prior work that normalized local brain measures by a whole-brain mean to reduce inter-individual differences in global values^39,50^. It furthermore accounts for potential global differences in the scale of the BOLD signal across scans. Mean normalized fluctuation amplitude was quantified for all participants (n = 1033) by dividing an individual’s fluctuation amplitude in each parcellated cortical region by the average fluctuation amplitude computed across all cortical regions. Notably, because whole-brain mean fluctuation amplitude declined across the age range studied, regional trajectories in this sensitivity analysis represent regional age-related decreases or increases relative to this global change.

The sixth and final sensitivity analysis was implemented to verify that the hierarchical sequence of fluctuation amplitude maturation would be observed when using a different atlas for cortical parcellation. As described above, vertex-wise fluctuation amplitude data in fsLR surface space was parcellated with the Schaefer-400 atlas. GAMs were fit using data from the main study sample of 1033 individuals for each Schaefer atlas region.

### Characterizing environmental effects

#### Neighborhood-level environment factor scores

Prior work suggests that social, economic, and physical features of children’s neighborhood environments can influence brain maturation and plasticity. To ascertain whether the developmental environment may specifically influence cortical activity dynamics, we studied associations between spontaneous BOLD fluctuations and neighborhood-level environment factor scores; the derivation of these factor scores has been previously explained in detail^55^. Briefly, geocoded information about each individual’s neighborhood environment was extracted using their home address and the census-based American Community Survey. The first factor from an exploratory factor analysis conducted on census variables by Moore et al. (2016)^55^ is used in the present study. This neighborhood-level environment factor score had positive loadings for the percent of residents who are married (loading = 0.85), median family income (0.82), the percent of residents with a high school education (0.74), the percent of residents who are employed (0.68), and median age (0.61) as well as negative loadings for the percent of residents in poverty (−0.86), population density (−0.71), and the percent of houses that are vacant (−0.60), and a weak loading for the percent of residents who are female (−0.26).

#### Generalized additive models

GAMs were used to investigate whether variability in neighborhood environments was associated with variability in regional fluctuation amplitude. For each region in the HCP multimodal atlas, a GAM was fit with fluctuation amplitude as the dependent variable, age as a smooth term, and sex, in-scanner motion, and the environment factor score as covariates. The *t*-value associated with the factor score term in each GAM represents the magnitude and direction of the fluctuation amplitude-environment relationship.

#### Alignment with the sensorimotor-association axis

Variability in youth’s neighborhood-level environments may differentially affect intrinsic activity in areas of the brain with different cortical properties and developmental trajectories. To assess whether environment effects were systematically related to cortical organization and development, we tested for an association between environment factor score *t*-values and S-A axis ranks using a Spearman’s correlation.

### Spin-based spatial permutation testing

Cortical data often exhibit distance-dependent spatial autocorrelation that can inflate the significance of correlations between two cortical feature maps. To mitigate this issue, we assessed the significance of each Spearman’s correlation that compared two whole-brain cortical feature maps with spin-based spatial rotation tests, or “spin tests”^107,108^. Spin tests compute a *p*-value (denoted *p*_spin_) by comparing the empirically observed correlation to a null distribution of correlations obtained by randomly spatially iterating one of the two cortical feature maps. In particular, spin tests generate a null by rotating spherical projections of one feature map while maintaining its spatial covariance structure. Here, we generated a null distribution based on 10,000 spherical rotations. Spin tests were implemented using the *rotate_parcellation* algorithm in R^109^.

### Data and code availability statement

This paper analyzes existing, publicly available data from the Philadelphia Neurodevelopmental Cohort, accessible from the Database of Genotypes and Phenotypes (phs000607.v3.p2) at https://www.ncbi.nlm.nih.gov/projects/gap/cgi-bin/study.cgi?study_id=phs000607.v3.p2. All original code has been deposited at Zenodo and is available at https://doi.org/10.5281/zenodo.6959989. A detailed description of the code and guide to code implementation is additionally provided at https://pennlinc.github.io/spatiotemp_dev_plasticity/. Any additional information required to reanalyze the data reported in this paper is available from the lead contact upon request.

## ACKNOWLEDGEMENTS

This study was supported by grants from the National Institute of Health: R01MH113550 (TDS & DSB), R01MH120482 (TDS), R01MH112847 (RTS & TDS), R01MH119219 (RCG & REG), R01MH123563 (RTS), R01MH119185 (DRR), R01MH120174 (DRR), R01NS060910 (RTS), R01EB022573 (TDS), RF1MH116920 (TDS & DSB), RF1MH121867 (TDS), R37MH125829 (TDS), R34DA050297 (APM), K08MH120564 (AAB), K99MH127293 (BL), and T32MH014654 (JS). VJS was supported by a National Science Foundation Graduate Research Fellowship (DGE-1845298). The PNC was supported by MH089983 and MH089924. Additional support was provided by the Penn-CHOP Lifespan Brain Institute.

## AUTHOR CONTRIBUTIONS

Conceptualization: VJS, BL, TDS Methodology: VJS, BL

Software: VJS, BL, AA, MAB, MC, SC

Validation: BL Formal Analysis: VJS

Resources: REG, RCG, TDS

Data Curation: AA, MAB, MC, SC, TDS

Writing-Original Draft: VJS

Writing-Review and Editing: VJS, BL, JS, AA, AAB, DSB, MAB, MC, SC, YF, REG, RCG, APM, TMM, DRR, RTS, TDS

Visualization: VJS Supervision: TDS

Funding Acquisition: YF, REG, RCG, RTS, TDS

